# The WD Domain Of *Atg16l1* Crucial For LC3-Associated Phagocytosis Is Not Required For Preserving Skin Homeostasis In Mice

**DOI:** 10.1101/2023.10.30.564683

**Authors:** Shannon Conway, Matthew Jefferson, Derek T. Warren, Thomas Wileman, Christopher J. Morris

## Abstract

The skin is a multifunctional organ, forming a barrier between the external and internal environment, thereby functioning as an initial safeguard against foreign and harmful environmental and biological factors. The barrier function is governed by the epidermis, a stratified epithelium composed of an outermost cornified envelope, several differentiated layers, and an innermost basal layer, anchored to the basement membrane (BM). Below the BM lies the dermis, which is composed of an extracellular matrix (ECM), formed by the dermal fibroblasts and dermal adipose tissue. The skin is subject to constant cellular turnover (terminal differentiation) that maintains the homeostatic state of the skin. Recently, autophagy has been implicated in epidermal differentiation and in preserving homeostasis in the skin. The AKT/mTORC1 pathway is a key regulator for epidermal development and differentiation, and conditions (e.g., psoriasis) are linked to the hyperactivation of these key autophagy regulators. A further process known as LC3-associated phagocytosis (LAP) uses some but not all components of autophagy. The Atg16l1 E230 mouse model (E230), deficient for LAP, has been widely used to study the effects of LAP-deficiency and autophagy in tissue homeostasis.

Here, the E230 model was used to study the relationship between skin homeostasis and LAP and to determine whether LAP-deficient mice (WD) display a cutaneous skin phenotype. Histological analysis of male 1-year-old Wild Type (WT) and WD mice skin revealed morphological differences in dorsal and tail skin. qPCR analysis of key keratins showed no differences (p > 0.05) when compared between genotypes. This was confirmed by western blotting studies. In addition, the expression of general proliferation markers (Akt & ERK2) showed a small reduction (p < 0.05) in the WD mice skin. General skin barrier formation was assessed by dye permeation assays, which demonstrated full and proper formation of the skin barrier at E18.5 in both WT and WD. Notable dermal thinning in the WD mice skin (p < 0.0001), led us to examine the biomechanical properties to dismiss any abnormalities relating to biomechanical functionality. Results indicate that WT and WD mouse skin show identical biomechanical properties at these ages (p < 0.05 and p < 0.01, respectively). In summary, the noted differences in the dermal and epidermal histology of WD skin were shown to be functionally insignificant in terms of epidermal permeability and dermal biomechanics.

## Introduction

The skin is a multifunctional organ, acting as an initial safeguard against foreign and harmful environmental and biological factors (Joost et al., 2020). It undergoes constant dynamic renewal, where a variety of stem and progenitor cells are activated in its three unique layers – the epidermis, dermis, and hypodermis (Joost et al., 2020). Keratinocytes lost from the superficial epidermis through desquamation or injury are replenished by the terminal differentiation of cells originating from a stem cell compartment found at the dermal-epidermal junction (Joost et al., 2020). In recent years, a role for autophagy has been implicated during epidermal differentiation and in maintaining skin homeostasis (Rossiter et al., 2013; Akinduro et al., 2016). Autophagy plays a key role in maintaining tissue homeostasis by degrading damaged proteins and organelles in lysosomes (Rai et al., 2019). AKT and mTORC1, which inhibit autophagy, have been shown to be key regulators for epidermal development and differentiation (Akinduro et al., 2016). Conditions (e.g., psoriasis), have been linked to the hyperactivation of mTORC1, while inhibition of this key regulator favours the expression of essential autophagy markers (e.g., LC3) (Akinduro et al., 2016). A pathway related to autophagy called LC3-associated phagocytosis (LAP) is activated in response to recognition of phagocytic cargoes by Toll-like receptors (Rai et al., 2019). LAP uses a subset of autophagy proteins to attach LC3/ATG8 to the cytosolic side of the phagosome membrane to facilitate fusion with lysosomes, thereby degrading material entering the cells (e.g., pathogens and protein aggregates) (Rai et al., 2019). A LAP-deficient mouse model, known as Atg16l1^ΔWD^ lacks the WD domain of *Atg16l1*, which is required for LAP, but not autophagy (Supplementary Figure S1) (Rai et al., 2019). Atg16l1^ΔWD^ mice have been used to determine the roles played by autophagy and LAP in maintaining tissue homeostasis *in vivo* (Rai et al., 2019; Slowicka et al, 2019). LAP is reported to be critical in orchestrating mucosal defences to microbial infection, including influenza and *Listeria monocytogenes* (Wang et al., 2021; Gluschko et al., 2022). The Atg16l1^ΔWD^ mouse model also develops spontaneous Alzheimer’s disease, where the WD domain of *Atg16l1* is required for efficient clearance of β-amyloid and the prevention of neuroinflammation (Heckmann et al., 2020). Collectively, these observations indicate a major role for the *Atg16l1* WD domain in barrier function and in reducing inflammation and led us to hypothesise that Atg16l1^ΔWD^ mice would display a cutaneous phenotype.

## Results & Discussion

Here, the skin histology of 1 year-old male, wild-type (WT) and Atg16l1^ΔWD^ (i.e., ΔWD) littermate mice, housed under standard conditions, was assessed by H&E staining and the thickness of the individual skin strata was measured at various anatomical locations (Figure 1a; Table S3). In the 1-year-old WT and ΔWD mice the quality and quantity of the hair follicles remained unaffected, with a few white hairs present (Figure 1a-b; Figure S2). Thinning of the epidermis was observed in the dorsal skin, while the dermis showed consistent thinning across anatomical locations, despite the similar growth rates and body mass, observed in this model (Table S3) (Rai et al., 2019).

**Figure 1.**
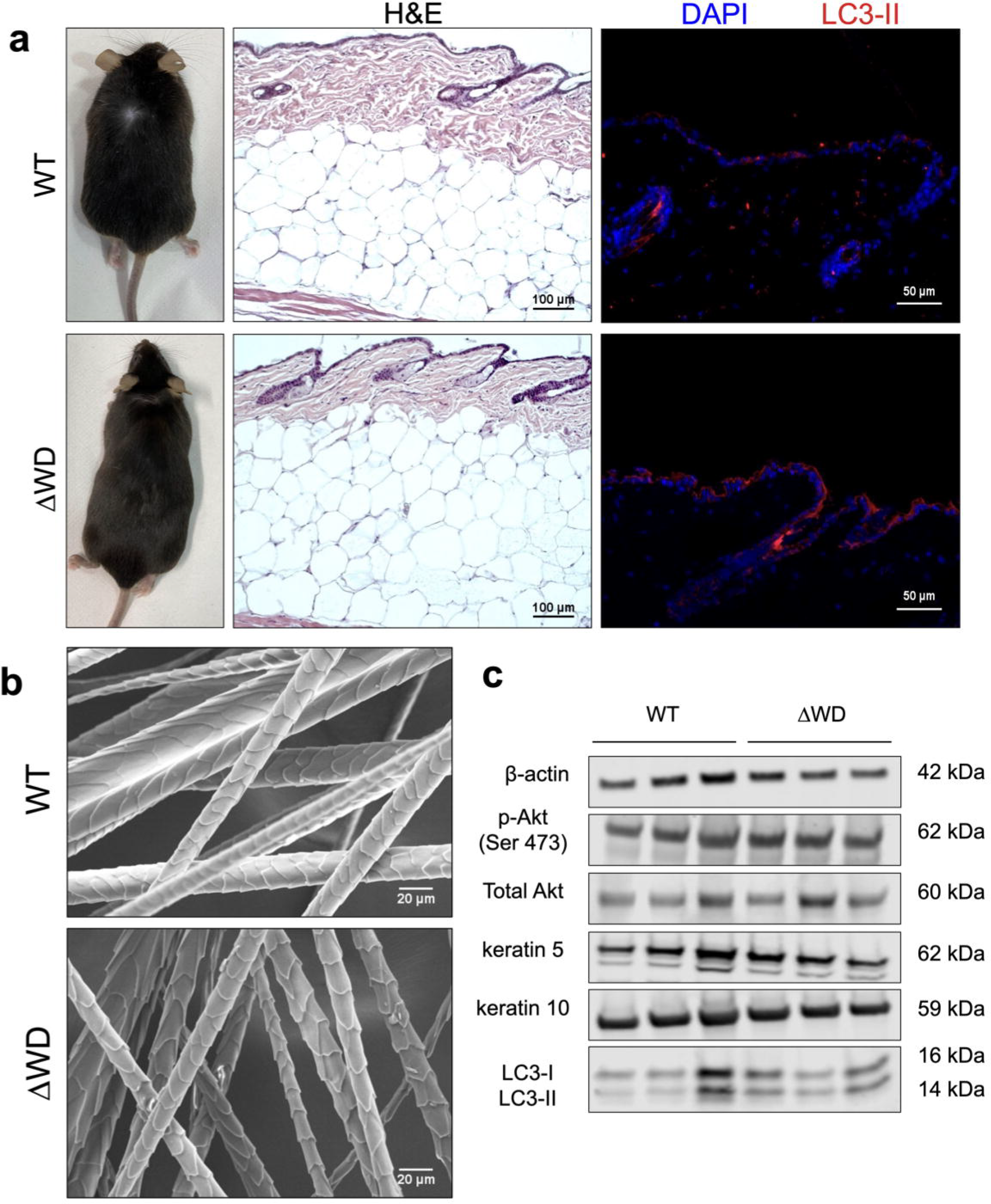
Histological analysis of 1-year-old Atg16l1^ΔWD^ mice showed epidermal and dermal thinning in ΔWD mice *cf*. WT littermate controls. **(a)** Overall appearance of WT and ΔWD mice. Histological examination of WT and ΔWD skin, showing epidermis, dermis, and hypodermis. Aggregated LC3-II expression (red) in ΔWD dorsal skin in comparison to WT littermate control. Scale bar: 100 μm (left) 50 μm (right). **(b)** Scanning electron microscopy (SEM) of hair follicles plucked from WT and ΔWD dorsal skin. Scale bar: 20 μm. **(c)** Western Blot of keratin 5, keratin 10, Akt, p-Akt and LC3-I/II. No changes in protein expression of keratins and LC3-I/II between WT and ΔWD (p > 0.05). Small reduction in expression of Akt in ΔWD *cf*. WT (p < 0.05). N=6, Unpaired t-test.

Epidermal thinning in the ΔWD mice prompted us to examine the mRNA and protein levels of key epidermal keratins and proliferative markers. Epidermal permeability and barrier function is closely linked to the precisely controlled turnover and replacement of epidermal cell layers (Akinduro et al., 2016; Harn et al., 2021). Indeed, dysregulation of such, results in skin diseases including psoriasis (epidermal thickening) and age-related skin thinning (Akinduro et al., 2016; Harn et al., 2021). qPCR analysis revealed no change in the basal layer marker, *KRT5* and upper epidermal markers, *KRT1, KRT10* and *IVL* between WT and ΔWD mice (Figure S3). Western blotting for keratin 5 and keratin 10 confirmed the RNA expression data (Figure 1c). Blotting for total and phosphorylated Akt showed there to be a small reduction in the expression of this proliferation marker in the ΔWD mice (Figure 1c).

This indicates that the WT and ΔWD mice display normal epidermal proliferation and differentiation. In addition, both the WT and ΔWD mice appear to generate consistent levels of lipidated LC3-II under steady-state conditions, highlighting that the ΔWD mice are still capable of generating LC3 in response to epidermal cellular turnover (Figure 1c). Nevertheless, LC3 in 1-year dorsal ΔWD skin appeared to aggregate throughout the epidermis compared to WT skin, indicating a potential lack of LC3 degradation in the ΔWD mouse (Figure 1a).

Intact skin permeability was assessed using dye permeation assays on E18.5 embryos. At this stage of development, the epidermal strata have fully developed, and the absence of the hair presents an uninterrupted epidermal surface barrier; the permeability of which can be assessed using toluidine blue (TB) and sodium fluorescein (SF) dyes (Figure 2a). No changes in the extent of TB retention in the skin was noted (Figure 2a); dye retention was noted only at the umbilicus and sites of tail cutting for genotyping. Quantification of SF dye penetration indicated a modestly reduced permeability of ΔWD skin compared to WT skin (Figure 2b). This indicates that the loss of LAP functionality has no detrimental effect on the barrier function of the epidermis.

**Figure 2.**
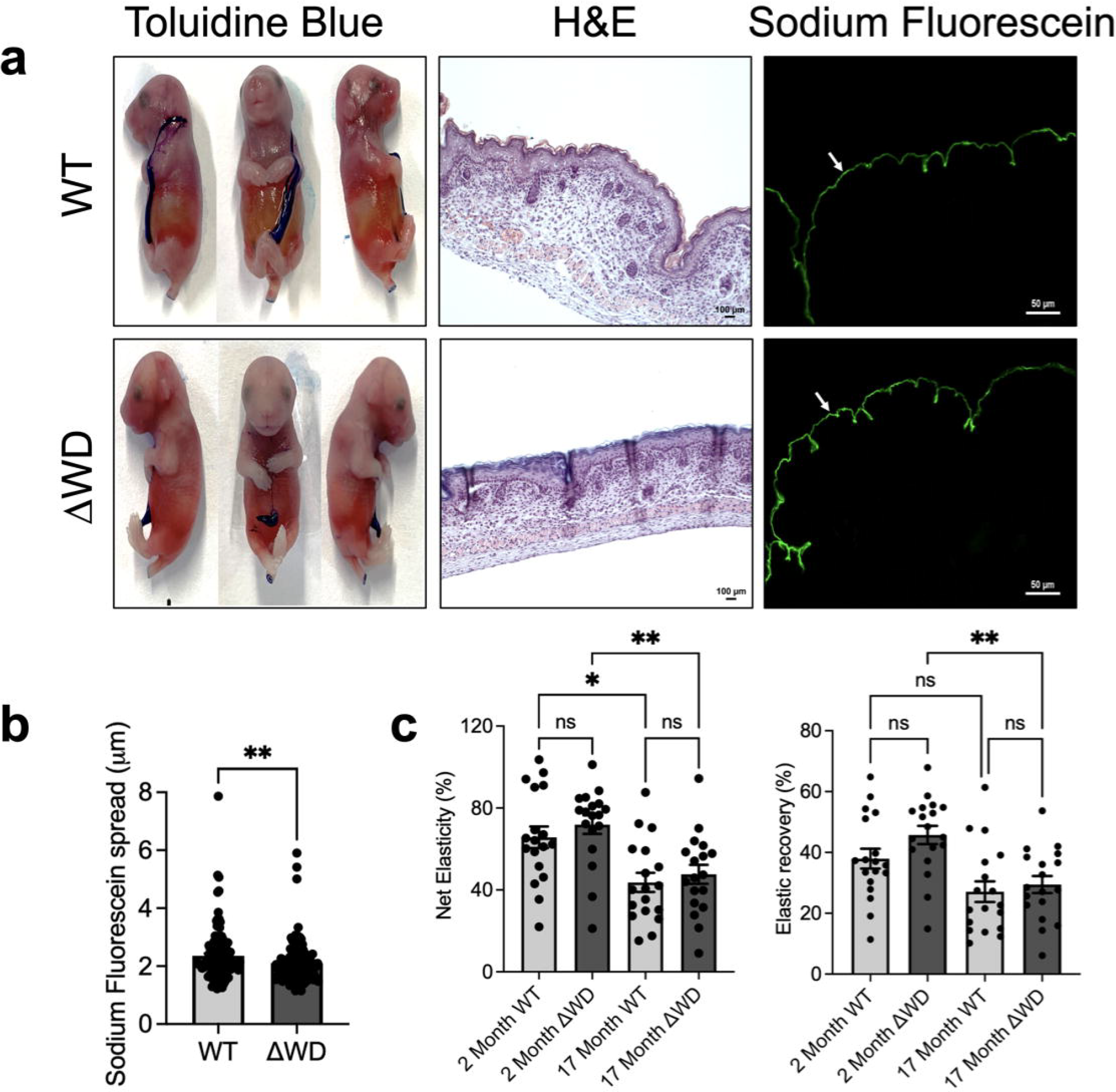
Assessment of the Atg16l1^Δ^WD mouse model skin barrier in E18.5 embryos revealed an intact skin barrier. **(a)** Left: Dye permeation assay on E18.5 embryos using toluidine blue dye; middle: H&E images showing epidermis, dermis, and hypodermis. Scale bar: 100 μm Right: Sodium Fluorescein (SF) dye accumulating in upper epidermal layer – stratum corneum (arrow). Scale bar: 50 μm **(b)** SF penetration into the embryonic epidermis. Data shown are mean ± S.E. N= 3, n= 40. ** p < 0.01. Unpaired T-test **(c)** Net elasticity (left) and elastic recovery (ER) (right) of dorsal 2-month and 17-month-old WT and ΔWD mouse skin. Data shown are mean ± S.E. N= 3, n= 6. * p < 0.05; ** p < 0.01. Two-way ANOVA.

As is the case for humans, the mechanical and structural properties of murine skin change with age (Chen et al., 2017; Harn et al., 2021). The dermal layer provides mechanical and structural support to the skin and serves as a cushion for the blood and lymphatic perfusion circuits [9]. As dermal thinning was observed in the ΔWD mouse, we hypothesised that the gross biomechanical functionality may be compromised in the ΔWD mouse. To test this possibility, the biomechanical properties of WT and ΔWD mouse skin was assessed. We evaluated the elastic function of the mouse skin in ages of 2 and 17 months. Both the net elasticity and elastic recovery parameters of the skin were comparable at matched ages in both genotypes (Figure 2c). As expected, both parameters were decreased at 17-months, in comparison to 2-month-old mouse skin. Overall, these results indicated identical extrinsic skin ageing in both genotypes.

## Conclusion

In conclusion, the histological examination of the 1-year-old male ΔWD mouse skin revealed morphological differences, when compared to the WT. The differences were shown to be functionally insignificant in terms of epidermal permeability and dermal biomechanics. The deletion of the WD domain of *Atg16l1* displayed no clear skin phenotype in healthy skin, yet the cutaneous response to cellular offences (e.g., viral, or bacterial infection) in the ΔWD mice may exhibit defective responses (e.g., cytokine release), which remains an ongoing study in our laboratory.

## Data availability statement

No datasets were generated or analysed during the current study. Data related to this article will be made available by the authors, without undue reservation.

## Conflict of interest statement

The authors state no conflict of interest.

## Acknowledgements

This work was supported by a studentship for SC from the Faculty of Science at the University of East Anglia, Norwich, UK. We would like to thank Mr. Bertrand Lézé for his help with the scanning electron microscopy. In addition, we would like to extend our thanks to Enviroderm, for providing us with the Cutometer® MPA580 for our biomechanical work. Lastly, we would like to thank Dr James McColl for his support with the fluorescent microscopy.

## Author Contributions Statement

**Shannon Conway**: Conceptualisation, Methodology, Investigation, Formal Analysis, Writing – Original Draft, Writing – Review & Editing, Visualisation, Project Administration. **Matthew Jefferson:** Methodology, Resources, Writing-Review & Editing, Project Administration.

**Derek T. Warren:** Conceptualisation, Methodology, Resources, Writing - Review & Editing, Supervision.

**Thomas Wileman:** Conceptualisation, Methodology, Resources, Writing – Review & Editing, Supervision, Funding acquisition.

**Christopher J. Morris:** Conceptualisation, Methodology, Resources, Writing – Original Draft, Writing – Review & Editing, Supervision, Funding acquisition.

## Notes

### Competing Interest Statement

The authors have declared no competing interest.

### Summary of Updates

Article title revised.

## References

1. O. Akinduro, K. Sully, A. Patel, D. Robinson, A. Chikh, G. McPhail, et al., Constitutive autophagy and nucleophagy during epidermal differentiation, Journal of Investigative Dermatology.136 (2016) pp.1460–1470.

2. H. Chen, X. Wang, J. Han, Z. Fan, S. Sadia, R. Zhang, et al., AKT and its related molecular feature in aged mice skin, PLoS One. 12 (2017) e0178969.

3. A. Gluschko, A. Farid, M. Herb, D. Grumme, M. Krönke & M. Schramm, Macrophages target Listeria monocytogenes by two discrete non-canonical autophagy pathways, Autophagy. 18 (2022) pp.1090–1107.

4. H.I.C Harn, C.C Chen, S.P Wang, M. Lei & C.M Chuong, Tissue mechanics in haired murine skin: potential implications for skin aging, Frontiers in Cell and Developmental Biology. 9 (2021) 635340.

5. B. Heckmann, B.J.W Teubner, E. Boada-Romero, B. Tummers, C. Guy, P. Fitzgerald, et al., Nancanonical function of an autophagy protein prevents spontaneous Alzheimer’s disease, Sciences Advances. 6 (2020) eabb9036.

6. S. Joost, K. Annusver, T. Jacob, X. Sun, T. Dalessandri, U. Sivan, et al., The Molecular Anatomy of Mouse Skin during Hair Growth and Rest, Cell Stem Cell. 26 (2020) pp. 441–457, e7.

7. S. Rai, M. Arasteh, M. Jefferson, T. Pearson, Y. Wang, W. Zhany, et al., The ATG5-binding and coiled coil domains of ATG16L1 maintain autophagy and tissue homeostasis in mice independently of the WD domain required for LC3-associated phagocytosis, Autophagy. 15 (2019) pp. 599–612.

8. H. Rossiter, U. König, C. Barresi, M. Buchberger, M. Ghannadan, C.F. Zhang, et al., Epidermal keratinocytes form a functional skin barrier in the absence of Atg7 dependent autophagy, Journal of Dermatological Science. 71 (2013) pp. 67–75.

9. K. Slowicka, I. Serramito-Gómez, E. Boada-Romero, A. Martens, M. Sze, I. Petta, et al., Physical and functional interaction between A20 and ATG16L1-WD40 domain in the control of intestinal homeostasis, Nature Communications, 10 (2019) 1834.

10. Y. Wang, P. Sharma, M. Jefferson, W. Zhang, B. Bone, A. Kipar, et al., Non-canonical autophagy functions of ATG16L1 in epithelial cells limit lethal infection by influenza A virus, The EMBO Journal. 40 (2021) e105543.

