## Supplementary Figures for "The WD Domain Of *Atg16l1* Crucial For LC3-Associated Phagocytosis Is Not Required For Preserving Skin Homeostasis In Mice"

**Supplementary Materials and Methods**

1. **Mice**

All experiments were performed in accordance with UK Home Office guidelines and under the UK Animals (Scientific procedures) Act 1986. The generation of the ${Atg16l1}^{\Delta\mathrm{WD}}$ mice has been previously described by (Rai et al., 2019). Generation and breeding of mice for the genotype was approved by the University of East Anglia Animal Welfare and Ethical Review Body and performed under UK Home Office Project License 70/8232.

2. **Tissue preservation and processing**

Dissected mouse skin was placed in a histology cassette and fixed in 10 % neutral buffered formalin (NBF) (Sigma) overnight at room temperature. The following day, the tissue was washed, dehydrated through a graded series of ethanol solutions and paraffin wax embedded. This process enables the generation of a solid cuboidal block containing the skin tissue, which can be easily mounted onto a microtome (Microm, HM355S) and stored at 4°C for long-term storage. 5 $\mu$m tissue sections were cut using the microtome blades (MX35 Ultra, Thermofisher Scientific) and were carefully transferred onto coated glass slides (Thermofisher Scientific, Superfrost Plus) and left to dry overnight at room temperature. Slides were labelled and stored at 4°C until further processing.

3. **Haematoxylin and Eosin (H&E) stain**

Tissue sections were H&E stained by submerging the slides in Mayers haematoxylin (Thermofisher) and Eosin Y (Thermofisher) solutions. Once the sections have passed through each solution, they were then mounted using coverslips (VWR) and mounting medium (DPX, Thermofisher Scientific). Sections were examined under the brightfield Apotome 3 Imager.Z2 microscope.

4. **Morphometric measurements of skin layers**

Brightfield images of H&E sections were analysed using the straight-line measurement tool in ImageJ (1.48v). A total of 20 measurements were taken in each distinctive skin layer at various anatomical locations (i.e., dorsal, ventral & tail), across 3 WT and 3 $\Delta$WD mice. Results were visualised and analysed in GraphPad Prism. The Mann-Whitney and unpaired t-test were used, where a $p <$0.05 is considered statistically significant.

5. **Epidermal-Dermal separation by enzymatic digestion using trypsin**

The entire mouse tail was removed and placed in PBS/Betadine solution (1:1), followed by a wash in PBS and rinse in 70% ethanol. Once the tail was washed, it was stored in a kanamycin solution (50 *µ*g/mL in PBS) until further processing. Skin was removed from the tail bone and placed epidermal side down in a sterile petri dish. The hypodermal layer was carefully removed before it was placed into a new sterile petri dish. The tail skin was incubated dermal side down in ice-cold 0.25% trypsin without EDTA (Speciality Media) for 2 hours at 37 °C. After incubation, the epidermis was peeled off with curved forceps and stored accordingly (i.e., in RNA later or snap frozen) for further processing.

6. **RNA extraction**

Mouse tissue was removed from RNA later (Thermofisher Scientific) and cut into small pieces (approx. 2 mm) and transferred to a 2 mL Eppendorf safe-lock tube (Fisher Scientific), containing 600 *µ*l TRIzol (Invitrogen) and one Tungsten Carbide bead (Qiagen). The tubes were placed in a tissue lyser (Qiagen, 85300) and were shaken twice for 5 minutes at 30 Hz. The homogenate was later centrifuged at 14,000 *x g* for 10 minutes at 4°C. The supernatant was removed and added to a fresh centrifuge tube (Eppendorf) containing 200 *µ*l of chloroform (Fisher Scientific). The tube was vortexed for 15 seconds and incubated at room temperature for 3-5 minutes. Following incubation, the solution was centrifuged at 14,000 *x g* for 15 minutes at 4°C to separate the phases. The upper aqueous layer was removed and added to a fresh centrifuge tube containing 200 *µ*l of 95% ethanol. The solution was mixed before proceeding with total RNA extraction using the GenElute total RNA purification kit (Sigma, RNB100), following manufacturer’s instructions. RNA was quality controlled using the Nanodrop 2000 (Thermofisher Scientific).

7. **cDNA synthesis and SYBR green qPCR**

RNA was reverse transcribed to cDNA using random hexamers (50 *µ*M), 10 mM dNTPs (New England BioLabs), 5X first-strand buffer (Invitrogen), 0.1 M DTT (Invitrogen), SuperScript II RT (Invitrogen) and 1 ng - 5 *µ*g total RNA. For relative quantification of mRNA expression using the 5 genes of interest, tail epidermal cDNA was analysed via SYBR green qPCR (PCR Biosystems), following manufacturer’s instructions. Pre-designed KiCqStart SYBR Green primers were purchased from Sigma (Table S1). SYBR Green qPCR was performed using the QuantStudio™ 3 Real-Time PCR System, 96 well (Thermofisher Scientific), using a MicroAmp optical 96-well reaction plate (Applied Biosystems). The SYBR green qPCR amplification program used for the analysis is shown in Table S2. Relative quantification was calculated using the $2^{-\Delta\Delta}$^CT^ method using 18S as a reference gene.

**Supplementary table S1: KicqStart SYBR Green Primers and 18S Housekeeping gene details**

| **Gene** | **Context Length Sequence (bp)** | **Sequence (5’-3’)** |
| --- | --- | --- |
| *18S* | 22  21 | Forward : GCCGCTAGAGGTGAAATTCTTG  Reverse : CATTCTTGGCAAATGCTTTCG |
| *KRT5* | 21  20 | Forward : GTGATGTTGAAGAAGGATGTG Reverse : TTCATGAAGTTGATCTCGTC |
| *KRT10* | 20  21 | Forward : CAATCAGAAGAGCAAGGAAC Reverse : CAGTGATTTCAGACTTATGGC |
| *KRT1* | 22  20 | Forward : CTACCAAATGGAAATGTCTCAG Reverse : GTAAAAGGTCTCAGCTTCAG |
| *IVL* | 21  20 | Forward : CTGTGAGTTTGTTTGGTCTAC  Reverse : GAAAGCCCTTCTCTTGAATC |

**Supplementary table S2: qPCR amplification program**

| **Step** | **Cycle** | **Time (s)** | **Temperature (°C)** |
| --- | --- | --- | --- |
| **Polymerase Activation** | 1 | 120 | 50 |
|  | 1 | 600 | 95 |
| **Denature** | 40 | 15 | 95 |
| **Anneal** | 40 | 60 | 60 |
| **Melt Curve** | 1 | 15 | 65 |
|  | 1 | 60 | 60 |
| **Dissociation** | 1 | 1 | 95 |

8. **Western Blot**

Tail skin isolated from WT and $\Delta$WD mice was snap frozen in liquid nitrogen and stored at

–80 °C until further processing. The tail skin was homogenised in a pestle and mortar and lysed in urea buffer (6.5 M urea, 1 mM DTT; 50 mM Tris-HCL). 1.5 *µ*l and 5 *µ*l of total tail skin lysate was boiled at 95 °C in SDS sample loading buffer and loaded into Bolt™ 4%-12% Bis-Tris SDS-Page gels (Invitrogen). The gels were electrophoresed alongside the Precision Plus Dual Colour protein standards (Bio-Rad) in NuPAGE MES SDS running buffer for 30 minutes at a constant voltage of 200V. Proteins were transferred onto PVDF membrane (Sigma), using the Bio-Rad Trans-Blot cell at constant 200 mA for 1 hour. Non-phosphorylated protein detection was carried out on PVDF membranes that were blocked in 5% skimmed-milk in Tris-buffered saline (TBS) plus 0.1% (v/v) Tween 20 (TBS-T). Phosphorylated proteins were detected following membrane blocking in 5% BSA in phosphate-buffered saline (PBS). The following primary and secondary antibodies were used in the analysis: anti-cytokeratin 10 (1:1000) (Abcam, EP1607IHCY); cytokeratin 5 (1:2000) (Sigma, SAB4501651); LC3A/B (1:1000) (Cell Signaling, 4108S); Akt (pan) (1:1000) (Cell Signaling, C67E7 ), p-Akt (1:10000) (Proteintech, 66444-1-Ig ); $\beta$-actin (1:10000) (Sigma, A2228), IRDye 800CW (1:10000) (Li-Cor, 926-32210) and IRDye 680RD (1:10000) (Li-Cor, 926-68071) . $\beta$-actin was used as an internal control. Blots were imaged using the Odyssey CLX imaging system (Li-Cor). Densitometry was performed using ImageJ (1.48v).

9. **Immunohistochemistry on paraffin-embedded tissue**

Sections of 5 *µ*m thickness were cut from each paraffin block and were placed on charged slides (Superfrost Plus slides; Thermo Scientific) and dried overnight at room temperature. Sections were de-waxed and rehydrated, before undergoing heat-mediated antigen retrieval using 0.1 M citrate buffer pH 6.0 for 10 minutes at 95°C. Tissue was permeabilized using 0.5% Tween-20 solution for 15 minutes. Endogenous peroxidase blocking was performed using 3% hydrogen peroxidase in 100% methanol for 20 minutes. Non-specific background blocking was carried out using 10% normal goat serum in PBS at room temperature for 30 minutes. Primary antibodies were diluted in 10% normal goat serum in PBS at their desired concentrations and were incubated at 4°C overnight in a humidified chamber. Rabbit LC3B (1:1000) (Sigma, L8918) and goat anti-rabbit Alexa Fluor 546 (1:1000) (ThermoFisher, A-11035). Tissue sections were mounted with Vectashield with DAPI (Vector Laboratories). Slides were imaged using the Apotome 3 Imager.Z2. Fluorescence intensity was quantified using ImageJ (1.48v).

10. **Timed mating of WT and** $\boldsymbol{\Delta}$**WD E18.5 embryos**

All ${Atg16l1}^{\Delta\mathrm{WD}}$mice (C57BL/6) were sourced from the Biomedical Research Centre animal facility. For the timed pregnancy, we used the trio-mating method (one male with two females per genotype). Matings were set up the evening before the start of the dark cycle. Females were separated from the male mouse upon confirmation of a vaginal plug the following day. This is considered as gestation day 0.5 (E0.5). If no plug was observed the female mouse was re-housed with the same sire for a maximum of five days. Embryos were harvested and sacrificed by anesthetic overdose at E18.5, 1 day before birth to ensure the full development of the skin barrier, without the presence of hair follicles.

11. **Skin permeability assessment – Toluidine Blue & Sodium Fluorescein**

Toluidine blue assay was done on unfixed whole mouse embryos (E18.5). The embryos were isolated from the pregnant female at E18.5 and sacrificed by anesthetic overdose (Euthatal, Pentobarbital sodium, 200mg in 1ml) and immersion in ice cold PBS for 30 minutes. The embryos were passed through a chilled methanol gradient, taking 2 minutes per step. After passing the embryos through methanol they were then immersed in 0.1% toluidine blue (Sigma) solution in water for 1-2 minutes on ice. The embryos were de-stained in PBS (pH 7.4) until a dye pattern was visible (NB. If a barrier defect is present).

Sodium fluorescein assay was done on unfixed whole mouse embryo (E18.5) dorsal skin. Embryos were placed epidermal side down in a petri dish containing 1 mM sodium fluorescein in PBS (pH 7.4) at 37 °C for 1 hour. Following 1 hour incubation, the dorsal skin was removed and fixed in 10% neutral buffered formalin (Sigma). After fixation, the tissue was dehydrated and processed as seen previously in section 2 – Tissue preservation and processing. Sodium fluorescein slides were rehydrated and imaged with the Zeiss ApoTome 2. Fluorescent dye uptake was quantified using ImageJ (1.48v).

12. **Elasticity measurement**

The Cutometer® MPA580 (EnviroDerm) with a 2-mm diameter probe was used in the assessment of the elasticity of the WT and $\Delta$WD mouse skin. The pressure in the probe was set to 400 mBar with 5 seconds of suction, followed by 5 seconds of release. To examine the elasticity in further detail, the following parameters were considered: net elasticity (Ur/Ue) and elastic recovery (Ur/Uf). Parameters were calculated in the MPA CTplus software.

13. **Scanning Electron Microscopy (SEM)**

Dorsal hair was plucked from the WT and $\Delta$WD mouse skin and mounted on aluminum stubs covered with conductive carbon tape and then gold coated with a Polaron SC7640 high resolution sputter coater, manufactured by Quorum Technologies. The hair samples were imaged under vacuum with the Zeiss Gemini 300 scanning electron microscope using the secondary electron (SE) detector. Hair follicle analysis was conducted using ImageJ (1.48v).

14. **Statistical Analysis**

| **Anatomical Location** | **Genotype & Age** | | | **p value** |
| --- | --- | --- | --- | --- |
|  | **Skin layer** | **WT 1 YR** | **𝝙WD 1 YR** |  |
| **Dorsal** | **Epidermis** | 13.54 ± 0.48 | 11.06 ± 0.50 | 0.0003 |
|  | **Dermis** | 251.90 ± 6.69 | 184.10 ± 7.41 | < 0.0001 |
| **Ventral** | **Epidermis** | 11.05 ± 0.54 | 10.38 ± 0.40 | 0.8929 |
|  | **Dermis** | 163.80 ± 5.97 | 111.10 ± 2.27 | < 0.0001 |
| **Tail** | **Epidermis** | 10.76 ± 0.36 | 10.14 ± 0.45 | 0.2897 |
|  | **Dermis** | 206.80 ± 4.34 | 120.90 ± 2.65 | < 0.0001 |

Data are expressed as means $\pm$ standard error of the mean. Normally distributed data were analysed with the unpaired t-test, while not normally distributed data was analysed with the Mann-Whitney test. For comparison between multiple groups the one-way and two-way ANOVA were used. p > 0.05 (ns); * p < 0.05; ** p < 0.01 and *** p < 0.001.

Supplementary Table 3 – Overview of mean skin thickness (*µ*m) ± S.E

Supplementary Figures

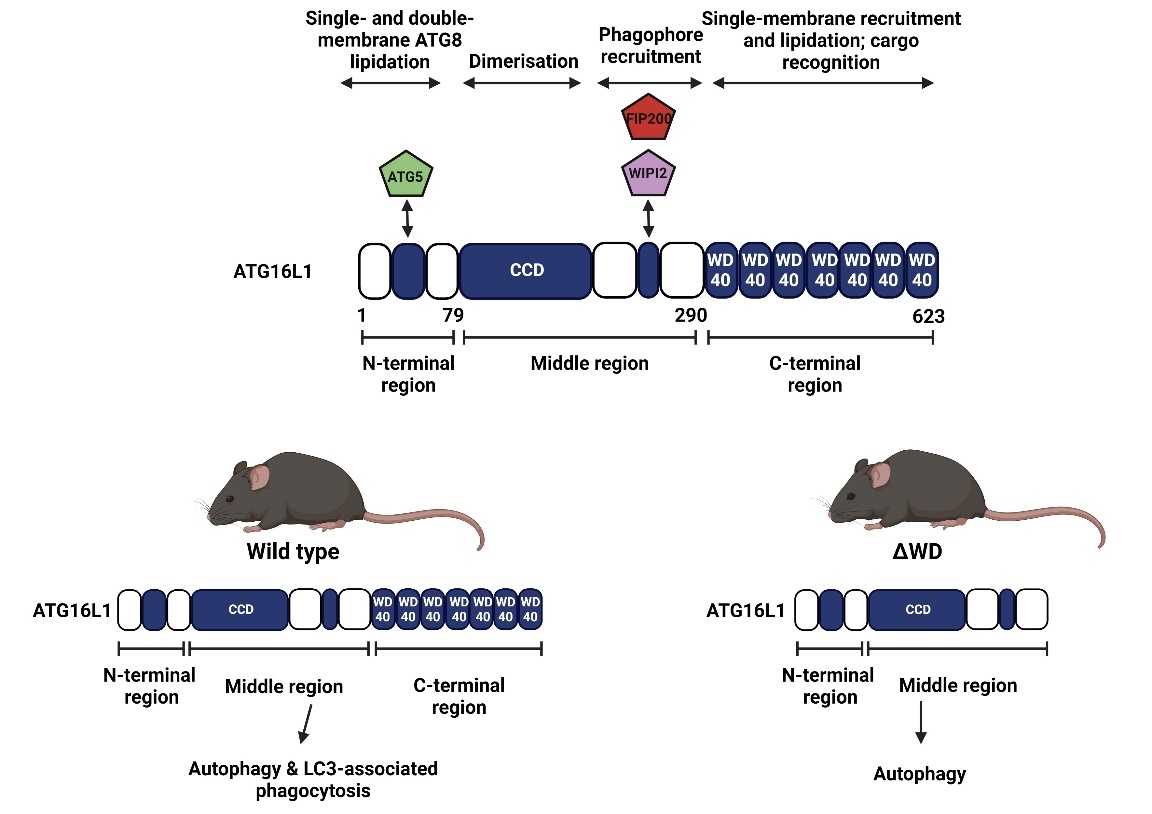

**Suppl. Fig. S1 Overview of Atg16l1^∆WD^ mouse model, showing removal of the WD domain leading to LC3-assciated phagocytosis deficiency.** Figure created with BioRender.com.

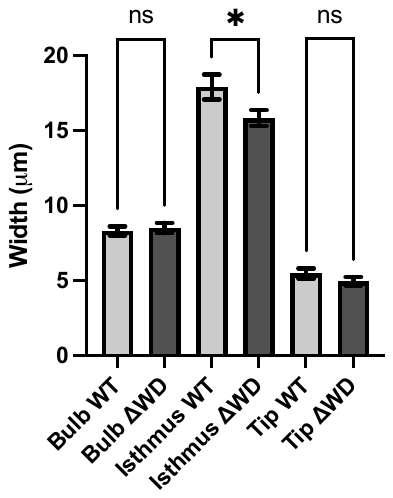

**Suppl. Fig. S2 Analysis of hair width at three distinct hair follicular regions.**

Measurements taken at bulbar region, isthmus and tip. Data shown are mean (n=30) $\pm$ S.E. Ordinary one-way ANOVA, * *p* < 0.05 and ns *p* > 0.05.

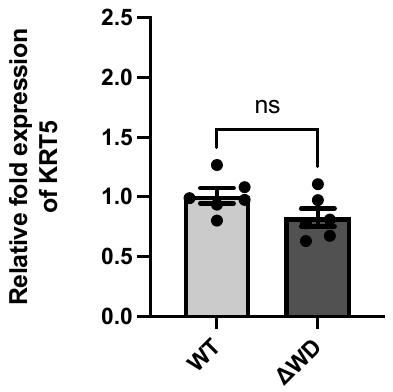

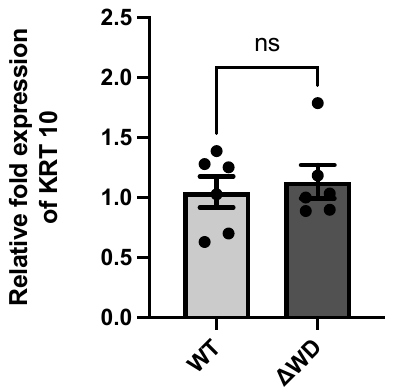

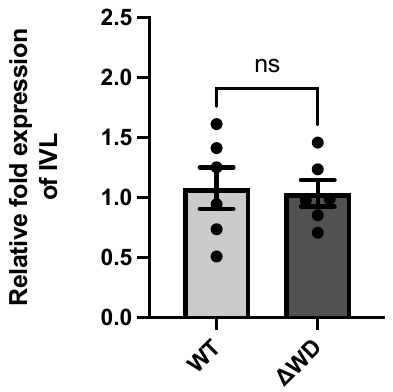

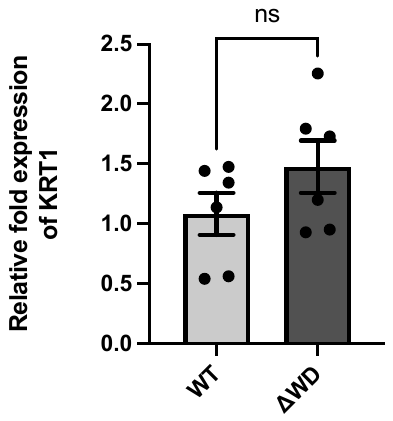

**Suppl. Fig. S3 qPCR analysis of key keratins (keratin 5, 10 & 1) and differentiation marker involucrin.** Relative mRNA expression normalized to 18S. Data shown are mean ± S.E. N=6, n=3. Unpaired t-test for normally distributed data (keratin 5, 1). Mann-Whitney test for not normally distributed data (keratin 10, involucrin). Ns p > 0.05.
